# Trace DNA from kill sites identifies predating tigers

**DOI:** 10.1101/2024.04.11.589017

**Authors:** Himanshu Chhattani, Abishek Harihar, Rounak Dean, Ajay Yadav, Kaushal Patel, Divyashree Rana, Awadhesh Pandit, Sanjay Kumar Shukla, Vincent Rahim, Uma Ramakrishnan

## Abstract

Predation ecology and evidence-based conflict management strategies require reliable and accurate identification of individual predators. Identifying predators is, however, complex, as they are secretive and individual identification is difficult. Trace DNA that predators leave behind at kill sites might provide an effective strategy to identify them but remains poorly evaluated at scale. We use non-invasive genetic samples from kill sites to assess their utility for predator identification. We systematically investigated 198 livestock kills in two critical source tiger populations in central India: Kanha and Bandhavgarh Tiger Reserves. We collected 342 salivary swabs from carcasses, 33 scat and 395 shed hair samples as potential sources of predator DNA, and individual tigers were identified using up to 123 SNP markers. All three sample sources identified predator species with high success (>95%). We identified individuals (with at least one sample per kill site, based on >40 SNPs) at 86% of all kill sites where tigers were detected. Shed hair samples were most effective for individual identification, followed by saliva and scat. Sample source and sampling season were the primary determinants of the number of SNPs typed per sample and the success of individual identification. Based on the site and type of sample collection, we classify species and individuals into three categories: true predator (high confidence as predator), circumstantial predator (medium confidence) and predator uncertain (low confidence). Individuals were classified as a true predator at 72 sites, circumstantial predator at 34 sites and predator uncertain at 49 sites. Our protocol allowed us to differentiate between predators and scavengers, even when multiple tigers were detected at the same kill site. Surprisingly, ∼40% of Bandhavgarh’s tigers were identified at at least one kill site. We suggest that when paired with systematic kill site investigation and sample collection, these methods can be effectively used to understand predation ecology better and facilitate evidence-based conflict management.

## 1. Introduction

Variation in predation behaviour within a population can be significantly influenced by specific traits, such as sex, age, morphology/phenotype as well as individual behavioural specialisation (Berezowska-Cnota et al., 2023; Bolnick et al., 2003; Dickman & Newsome, 2015; Estes et al., 2003; Scholz et al., 2020; Voigt et al., 2018). Understanding these processes at the individual level is crucial but challenging because of difficulties in reliably establishing which individuals are involved in predation events. Large carnivores often range over vast home ranges, have overlapping territories (both with individuals of the same species as well as other carnivore species), exhibit sociality and engage in scavenging and kleptoparasitism (Balme et al., 2017; Chundawat et al., 2016; Périquet et al., 2015). These traits and their elusive nature and absence of distinct individual markings in some species make it difficult to attribute a predation event to a specific individual.

Human carnivore conflicts, such as large carnivores’ attacks on livestock and humans, present another case where reliable predator identification is critical (Goodrich, 2010; Swan et al., 2017; Treves & Karanth, 2003). Globally, conflict mitigation involves the removal of individuals through translocation or lethal measures (Linnell et al., 1999; Swan et al., 2017). However, a challenge with these approaches is unreliable identification and the consequent removal of non-target individuals (Treves & Karanth, 2003). This can not only leave conflict unsolved but can also cause social disruptions in carnivore communities, increase conflict frequency, and invite further public criticism (Athreya et al., 2011; Sinha, 2018; Woodroffe & Frank, 2005). Therefore, for such management interventions to be effective, it is important to make evidence-based decisions starting with reliable individual identification.

Predation events are rarely witnessed, and accurate identification during such events is even more arduous. Conventionally used methods, such as visual analysis of prey remains and deployment of camera traps at kill sites, can lead to unreliable identification. Field assessments are restricted to identifying species, are highly dependent on the field expertise of the examiner, and limit retrospective use of data (Mumma et al., 2014). They can suffer from inaccuracy when species overlap in killing and feeding characteristics (Verzuh et al., 2018). Camera traps can be highly efficient in identification if deployed at a predation site before a predation event, such as when monitoring bird nests (Steffens et al. 2012); however, their use is limited in post-predation events where they might capture visiting, scavenging species/individuals (Steffens et al., 2012). Further, identifying individual predators using images becomes a challenge when individuals lack unique natural marks for identification.

Trace DNA of the predator left at the kill sites in the form of its saliva, shed hair, urine, scat, etc., can often be the only evidence for conclusive predator identification. Non-invasive genetic sampling, therefore, presents a promising approach to reliable and accurate identification of predators from kill sites (Blejwas et al., 2006; Fotedar et al., 2019; Nichols et al., 2012; Sundqvist et al., 2008). This approach has advantages over conventional methods as it is more sensitive and safer (for researchers and animals), can distinguish individuals of a species even without natural markings, and, most importantly, can distinguish a predator from a scavenger when systematically sampled. While being practical, the success of genetic samples can strongly be influenced by factors such as abiotic conditions (e.g., temperature, rainfall, light), biotic factors (e.g., microbial activity, prey and predator species, maggot infestation), as well as sample collection procedures and storage methods (Harms et al., 2015; Nakamura et al., 2017; Piaggio et al., 2020; Reddy et al., 2012). If not collected systematically, genetic samples may also lead to incorrect attribution of predation events to a scavenger.

In this study, we attempt to evaluate and use non-invasive genetic samples from kill sites for predator identification. We do this by systematically sampling suspected tiger kill sites at two source populations in the central Indian tiger landscape. By collecting saliva, shed hair and scat samples of potential predators, we identify individual tigers using next-generation sequencing methods. We chose to sample livestock kills in these tiger reserves to evaluate molecular techniques for the following reasons: (i) livestock kills are frequently reported for financial compensation to the forest department, thereby providing opportunities to collect samples in comparison to wild prey kills, which are challenging to detect, (ii) livestock kills are quickly reported (usually <24 - 48 hours) providing an opportunity to collect fresh samples, (iii) multiple sample sources (saliva, shed hair, scat) can be collected from kill sites for comparison and, (iv) importantly, predator identification from livestock kill sites has management importance.

Specifically, we examined the influence of environmental factors such as season and kill site conditions on the number of SNPs typed and individual identification success. Based on our results, we propose a framework to identify and classify individual predators identified from a kill site and recommend sampling strategies for individual identification. Finally, we discuss the significance of our results, both methodological and conceptual, for conservation and management.

## 2. Materials and Methods

Fieldwork for this study was carried out in two phases. The first phase was conducted in Kanha Tiger Reserve (hereafter, Kanha) in 2017, during which we explored the availability and abundance of genetic samples along with varied approaches to kill site investigation, including sample collection. In the second phase, we conducted extensive sampling in Bandhavgarh Tiger Reserve (hereafter, Bandhavgarh) from April 2021 to March 2022. In both reserves, we systematically investigated livestock kills reported to be made by tigers by the livestock owners and forest department staff. We collected saliva, shed hair and scat samples as potential sources of predator DNA.

### 2.1 Study area

Kanha and Bandhavgarh have been classified as tropical moist deciduous forests with four main habitat types: grasslands, pure sal forest, miscellaneous forest and bamboo mixed forest (Awasthi et al., 2016; Champion & Seth, 1968). Kanha has an area of 2,074 km^2^ and is divided into two management units: the inviolate core (940 km^2^) with minimum human presence and a multiple-use buffer (1134 km^2^). Bandhavgarh spans 1,537 km^2,^ with a core of 717 km^2^ and a buffer of 820 km^2^. The reserves harbour over 100 tigers each (Kanha: 129, Bandhavgarh: 165) and are critical source populations for tigers in the central Indian landscape (Qureshi et al., 2023). The local economy is largely dependent on agriculture, livestock rearing and tourism. Kanha reserve has an estimated population of 137,600 humans along with 91,900 cattle in the core and buffer area (Negi & Shukla, 2011).

Bandhavgarh has at least 130 villages in the park’s buffer area with ∼110,000 cattle heads. The resulting overlap of high human use area with that of high tiger density often results in conflicts mainly in the form of livestock depredation (annual livestock kills in Kanha range from 400-600 (Negi & Shukla, 2011), while in Bandhavgarh range from 2500-2800) and, sometimes, human and tiger mortalities. Addressing conflict, therefore, is a key imperative for reserve management.

## FIELD METHODS

### 2.2 Kill site information

We investigated livestock kill sites that were reported by the livestock owners to the forest department for financial compensation. Both reserves have a similar, established system of reporting kills. When a livestock carcass is located and reported to the beat guard of the area by the livestock owner, the guard then verifies the claim by physically visiting the kill site and confirming the involvement of a carnivore in making the kill. Predator identification by the forest department from the kill site is based on field signs. Once verified, the beat guard passes the message to the concerned range office, which then shares the information with the park field director’s office to initiate compensatory payment. This process, from initial detection by the livestock owner to final reporting to the field director’s office, usually takes 24 to 48 hours.

At each kill site, we first noted the GPS location, date and time of sample collection, and identity of the investigating team and the sample collector. We collected information on prey species, age and kill date, time and rainfall from the livestock owner. We recorded the forest department’s assessment of the predating species. This was done before our investigation to avoid influencing the department’s assessment. Additional observations such as drag distance (in metres, from the inferred kill location to the location of the carcass, wherever visible), percentage consumption (based on visual estimate), feeding pattern (parts consumed, removal of tail and intestines, etc.), maggot infestation (visual estimate, categorised as high/medium/low), canopy cover over the carcass (visual estimate in %) were noted. Photographs of the carcass were taken at the site.

At each kill site, we began our examination by conducting a quick ‘walk through’ to gain an overview, identify sites with evidence and inferred kill location. An intensive search for evidence was carried out along the drag marks between the carcass location and the inferred kill location. We scanned animal trails and potential carnivore resting areas for shed hair and faecal (scat) samples. Our search around the kill site and on the trails was limited to a distance of 30 meters. This was done to reduce the chances of encountering other carnivore samples since both reserves are high carnivore density areas. Carnivore resting areas were identified based on anomalous vegetation and impressions on soil.

Pugmarks, scrape marks and scats detected near the kill sites were assigned to carnivore species based on characteristics like shape, size, etc., following published field manuals (Karanth & Nichols, 2002). Active care was taken not to disturb evidence during this search. However, the forest department/livestock owner had, in some cases, cleared the surrounding vegetation to ease access to the carcass before our arrival. Two people conducted searches independently to maximise sample collections and reduce detection bias. Sites where carnivore pugmarks of different characteristics (shape, size), multiple resting sites (of varying sizes), irregular lick marks and multiple scat samples were found, were noted as kill sites with multiple carnivore presence. Once initial scene documentation was completed, we decided on the collection order and initiated sample collection. We collected saliva samples first (starting from predation wounds) and then scat and shed hair to reduce the chances of sample contamination. We tried to decrease the time spent and disturbance caused at a kill site so as not to disturb the kill site.

### 2.3 Non-invasive genetic sampling

We collected genetic evidence from salivary marks, shed hair, and scats at each kill site. Sampling kits were not opened until all ancillary information was collected. To minimise the risk of on-field sample contamination, we changed gloves while collecting different samples and stored samples in different zip lock bags. Shed hair samples were collected independently from the various locations using sterile forceps and stored separately in zip lock bags or sterile 2 ml tubes to avoid sample contamination. Traces of potential carnivore saliva from the carcass and scat samples found near the kill site were collected using sterile polyester swabs. Efforts were made to collect at least three saliva swabs (one each from different parts of the carcass with at least one swab from the predation bite marks whenever available), multiple shed hair samples and all the scat samples detected at the kill site. The number and type of samples collected from each site also varied based on kill site conditions such as the presence of carnivores, scavengers and proximity to villages.

For swab collections, swabs were first briefly soaked in Longmire lysis buffer (Longmire et al., 1997) and then rolled over the target area for 10-12 seconds following Mumma et al., 2014. For scats, swabs were rolled over the outer, shiny areas preferred for host DNA. Samples were categorised as either predation or post-predation samples (Figure 1). Predation samples were samples that we suspected to be deposited during a predation event. These included saliva samples collected from predation wounds (distinguished from other bite marks based on the occurrence of haemorrhage) and shed hair samples collected from the inferred kill location (where the predation event occurred). Post-predation samples included saliva samples from lick and feeding marks, shed hair samples from suspected resting sites and sites near the carcass. Scat and uncategorised samples were categorised as post-predation samples. Swab samples were stored in 2 ml tubes with 1.5 ml Longmire lysis buffer.

**Figure 1:**
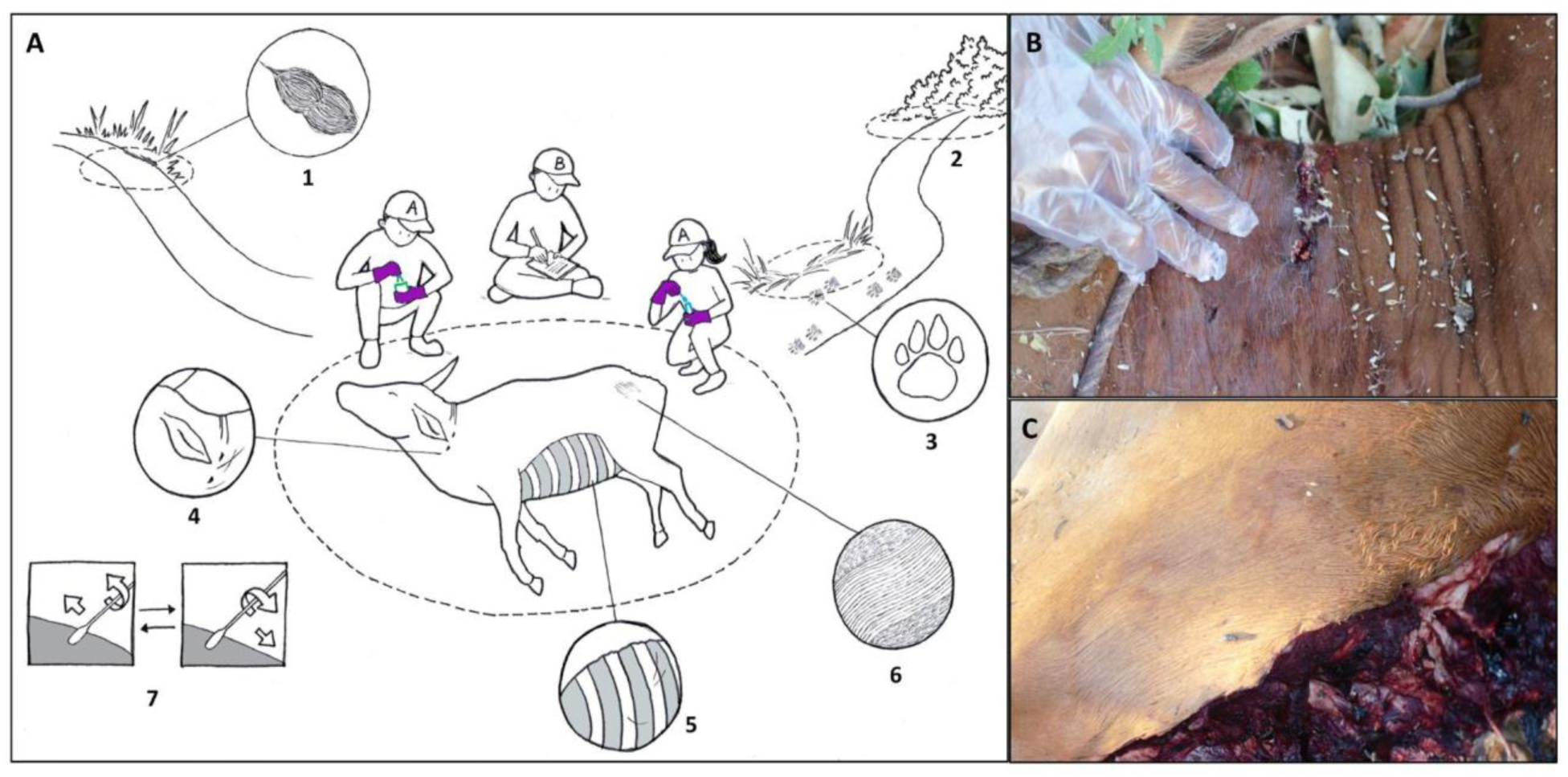
A) Field sampling scheme illustrating various types of genetic samples of potential predators collected at a kill site: 1) scat sample collected using swab 2) inferred kill location is a source of shed hair sample 3) resting sites are a source of shed hair samples, 4) predation marks in the neck region are a source of saliva samples, 5) feeding area source of saliva and shed hair samples, 6) lick marks characterised by unidirectional bend of hair are source of saliva samples and 7) showing swabbing technique. Samples collected from 2 and 4 were classified as predation samples, whereas the rest were classified as post-predation samples. B) Photograph of predation wound in neck region with visible saliva deposit around the puncture region (darker colour) and C) Photograph of carnivore lick areas with saliva deposit.

**Figure 2:**
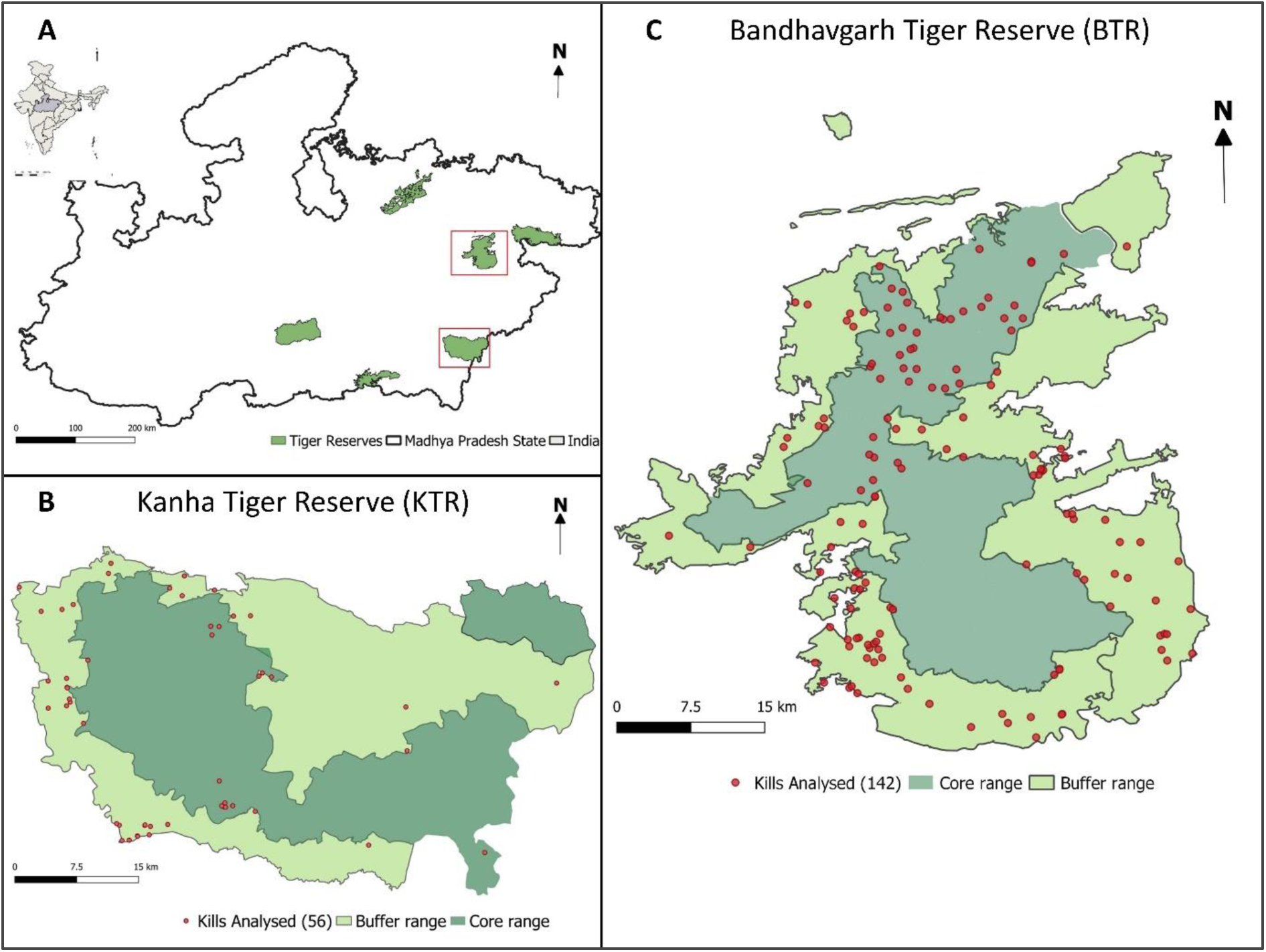
Study area map with sampling locations. A) Boundary of Madhya Pradesh state in India with study sites highlighted in red box. B) Kills sampled in Kanha Tiger Reserve (N = 56) and, C) Kills sampled inside Bandhavgarh tiger reserve (N = 142).

## GENETIC ANALYSIS

### 2.4 DNA extraction and species identification

DNA extraction was done in a dedicated low-concentration DNA extraction facility at the National Centre for Biological Sciences (NCBS), Bangalore. DNA extraction was done using QIAGen^®^ DNA Tissue and Blood extraction kits, following the protocol suggested by the manufacturer with modifications. Extraction controls (negatives) were set to detect any contamination. DNA extraction of different sample types was done separately to avoid contamination. All samples were screened using tiger-specific primers to identify positives (Bhagavatula & Singh, 2006). PCR controls were set to detect contamination while setting up the PCR reaction.

### 2.5 Individual identification

Samples genetically confirmed to be tigers were genotyped using a panel of 123 SNPs, as Natesh et al. (2019) described. This mainly involved a three-step process – a multiplex PCR, an indexing PCR, and library preparation and sequencing. Multiplex PCR of Kanha samples was performed using a unified pool of 123 primers. Primers were split into two pools (based on amplicon size) for Bandhavgarh samples. Indexing PCR was performed to add a unique combination of i5 and i7 Illumina indexes to samples. The indexed products were pooled and bead-purified to retain only amplicons of targeted size. The resulting library was then sequenced on an Illumina Miseq platform to obtain 75x2 paired-end reads. All the samples were processed in replicates.

Data filtering and subsequent individual identification were done independently for both study sites. Raw reads were filtered using the program TrimGalore (http://www.bioinformatics.babraham.ac.uk/projects/trim_galore/) with a quality value of 30 on phred33 scale to remove adapters and short reads. Filtered reads were mapped to the tiger genome assembled in the lab (Armstrong et al., 2021) using bwa mem (Li, 2013) with a penalty value of 3. Variant calling was done using bcftools (Li, 2011). Variants were filtered to retain only the targeted loci, following which genotype quality (10 or above) and depth (10 or above) filters using GATK (DePristo et al., 2011). Missing data filters were applied to remove loci missing in >15% of samples and samples with less than 40 SNPs.

After filtering, we retained 95 and 101 SNPs for Bandhavgarh and Kanha, respectively. Individuals were identified based on PI-HAT values using Plink (Purcell et al., 2007). PI-HAT values of known replicate pairs were used to determine the cut-off for recapture identification, as suggested by Natesh et al., 2019 and Sagar et al., 2021. Data of known replicates were merged to reduce missingness and re-analysed to obtain pairwise relatedness between unique samples after determining the relatedness cut-off.

Samples with pairwise relatedness greater than the recapture cut-off were identified as the same individual. Unique samples that had pairwise relatedness greater than 0.5 and lesser than the recapture cut-off with any sample were categorised as uncertain and discarded to avoid ambiguity. All other samples were identified as unique individuals without recaptures. Individual identification was done using Program R v 4.0.2.

### 2.6 Predicting SNP typing and individual identification success

We conducted standard Generalised Linear Models (GLM) analysis using Program R v 4.0.2 to assess how season and sample type influence the number of SNPs typed and individual identification success. We also evaluated the impact of sampling lag (number of days from predation event till sampling) and canopy cover (in percentage), but they were excluded from the final model as there was insufficient coverage. Rainfall and maggot infestation showed association with season and were therefore excluded. We used only Bandhavgarh samples to develop our models since there weren’t enough samples from Kanha across the seasons, and the lab protocols for individual identification were different from those of Bandhavgarh. The SNP count used for the analysis was the number of SNPs retained following filtering for genotype quality and depth. Similarly, any sample having more than 40 SNPs following filtering (depth (10), quality (10) and removing missing SNPs) were classified as successful for individual identification.

### 2.7 Predator assignment

Based on the location and type of sample used for identification, we classified tigers into three categories: True predator, circumstantial predator, and predator uncertain. Each of these assignments was at both species and individual levels. True Predators are species, individuals that were solely identified using predation samples, i.e. saliva samples from predation wounds on the carcass and shed hair samples collected from the location of kill. Circumstantial predators are species and/or individuals that were likely to be predators but were not identified from the predation samples either because predation samples were not collected or were ineffective in identification. Circumstantial predators were identified only when multiple samples were collected from a site, and all resulted in the same identification. Any sample where species and/or individuals identified from a kill site that did not fit the above categories were assigned as predator uncertain (schematic flow chart in Supplementary Figure S1). The highest confidence was assigned to true predator, followed by circumstantial predator and the least for predator uncertain, when ranking the likelihood of a species or individual being involved in the predation event.

## 3. Results

### 3.1 Sample availability and species identification

We collected sampled 342 saliva swabs, 395 shed hair and 33 scat samples from 198 kill sites (Bandhavgarh: 142 and Kanha: 56) (Supplementary Table S1). Shed hair and saliva samples were the most abundant source of DNA and were collected from 85% and 72% of kill sites sampled respectively. Shed hair samples were the only source of DNA for 52 kill sites, while saliva samples were the only source of DNA for 27 sites. Scat samples were found at 10% of the kill sites. At least one predation sample (saliva from the predation wound or shed hair from the inferred kill location) was collected from 153 (78%) kill sites. Season influenced the availability of saliva samples from predation wound, with samples collected from fewer kill sites in monsoon (28%) compared to winter (60%) and summer (60%).

Tigers were detected at all kill sites with species identification success with 98.5% saliva swabs, 98.2% shed hair and 96.9% scat samples. Using tiger-specific primers limited our detections to tigers; therefore, no amplification doesn’t indicate the absence of carnivore DNA. Season did not significantly influence the species identification success of any sample types.

### 3.2 Impact of season and sample type on number of SNPs typed

Using saliva samples from the winter season as the reference group, our models indicate a 30% and 53% decline in SNPs typed in summer and monsoon, respectively (Table 1). For shed hair (again, hair in winter as reference), an 11% and 37% decline in SNPs typed was observed in summer and monsoon, respectively. Shed hair samples typed more SNPs in comparison to saliva samples across seasons. In winter, the number of SNPs typed was highest for saliva and shed hair samples. A stronger seasonal impact was observed for saliva samples than shed hair samples.

**Table 1:** Predator classification scheme with conditions for assignment.

| Classification | Level | Sample used for identification | Condition | Confidence Assignment |
| --- | --- | --- | --- | --- |
| True predator (TP) | Species | Predation | 1. Species identified using only saliva samples from predation marks or shed hair samples from inferred kill location. | High |
|  | Individual | Predation | 1. Individual identified using only saliva samples from predation marks or shed hair samples from inferred kill location. |  |
| Circumstantial predator (CP) | Species | Post-predation | 1. No predation mark/inferred kill location sample<br>2. All/multiple samples from the kill site should identify only one species from the kill site. | Moderate |
|  | Individual | Post-predation | 1. No predation mark/ inferred kill location sample for species identification.<br>2. Individual identified from samples at the kill site, but not<br>predation mark/ inferred kill location.<br>3. All/multiple samples identify a single individual from the kill site. |  |
| Predator uncertain | Species | Post-predation | 1. Criteria for either TP or CP cannot assign species.<br>2. More than one potential predator species is identified at the kill site using samples other than predation samples.<br>3. When the species has been identified from just one sample, another than the predation sample. | Low |
|  | Individual | Post-predation | 1. Individual cannot be identified using criteria for TP or CP.<br>2. More than one potential predator individual identified at the kill site using samples other than from predation mark/inferred kill location .<br>3. Has been identified from just one sample other than from the predation mark/ inferred kill location. |  |

### 3.3 Individual identification success

After applying missing data filters, we set 40 SNPs as a cut-off for individual identification. Data filtering and subsequent individual identification were conducted independently for each study site. Of 757 unique tiger-positive samples processed, 449 samples (59%) had more than 40 SNPs and were used for individual identification (Table 2). PI-HAT scores of known replicates ranged from 0.64 to 1 for Kanha and from 0.69 to 1 for Bandhavgarh (Supplementary Figure S2). A recapture cut-off of 0.81 was set for Kanha and Bandhavgarh based on the 97.5% percentile of the relatedness distribution (between replicates). Seventy-two tigers (Kanha 19 and Bandhavgarh 53) were identified with varying numbers of recaptures (pairwise relatedness of Bandhavgarh individuals in Supplementary Figure S3). Four individuals from Kanha and 11 from Bandhavgarh had no recaptures and were detected only once. Seventy-one samples were classified as uncertain as they didn’t have more than 0.81 relatedness with any sample and/or had greater than 0.55 relatedness with multiple individuals. These samples were discarded to avoid ambiguity.

**Table 2:** Results of Generalised Linear Models (GLM) examining the influence of sample type (saliva and shed hair) and season (winter, summer and monsoon) on the number of SNPs typed.

| Model | Estimate | Std. Error | Pr(>\|z\|) |
| --- | --- | --- | --- |
| Saliva * Winter<br>(intercept) | 3.71379 | 0.01469 | <2e-16*** |
| Saliva * Summer | -0.35052 | 0.02816 | <2e-16*** |
| Saliva * Monsoon | -0.75672 | 0.04481 | <2e-16*** |
| Shed Hair * Winter | 0.42914 | 0.01788 | <2e-16*** |
| Shed Hair * Summer | 0.23086 | 0.23086 | <2.52e-12*** |
| Shed Hair * Monsoon | 0.29070 | 0.04850 | 2.05e-09*** |

At least one tiger was identified from 86% of all kill sites sampled. Shed hair samples were the most successful source (82%) for individual identification across seasons, followed by saliva (59%) and scat (58%) at kill sites (Table 2). Shed hair and saliva samples were the sole source of individual identification at 80 and 29 sites, respectively. Predation samples identified individuals at 53% of the sites where they were collected. Notably, both types of predation samples (saliva associated with predation mark and shed hair associated with predation mark) identified the same individual at 10 of 11 kill sites.

Our models indicate a higher probability of individual identification with shed hair samples when compared to saliva samples. Using the winter season as the reference group, the probability of individual identification declined by 10% and 30% in summer and monsoon, respectively, for saliva samples (Table 3). Similarly, for shed hair samples, the probability of individual identification decreases by 21% and 34% in summer and monsoon compared to winter.

**Table 3:**
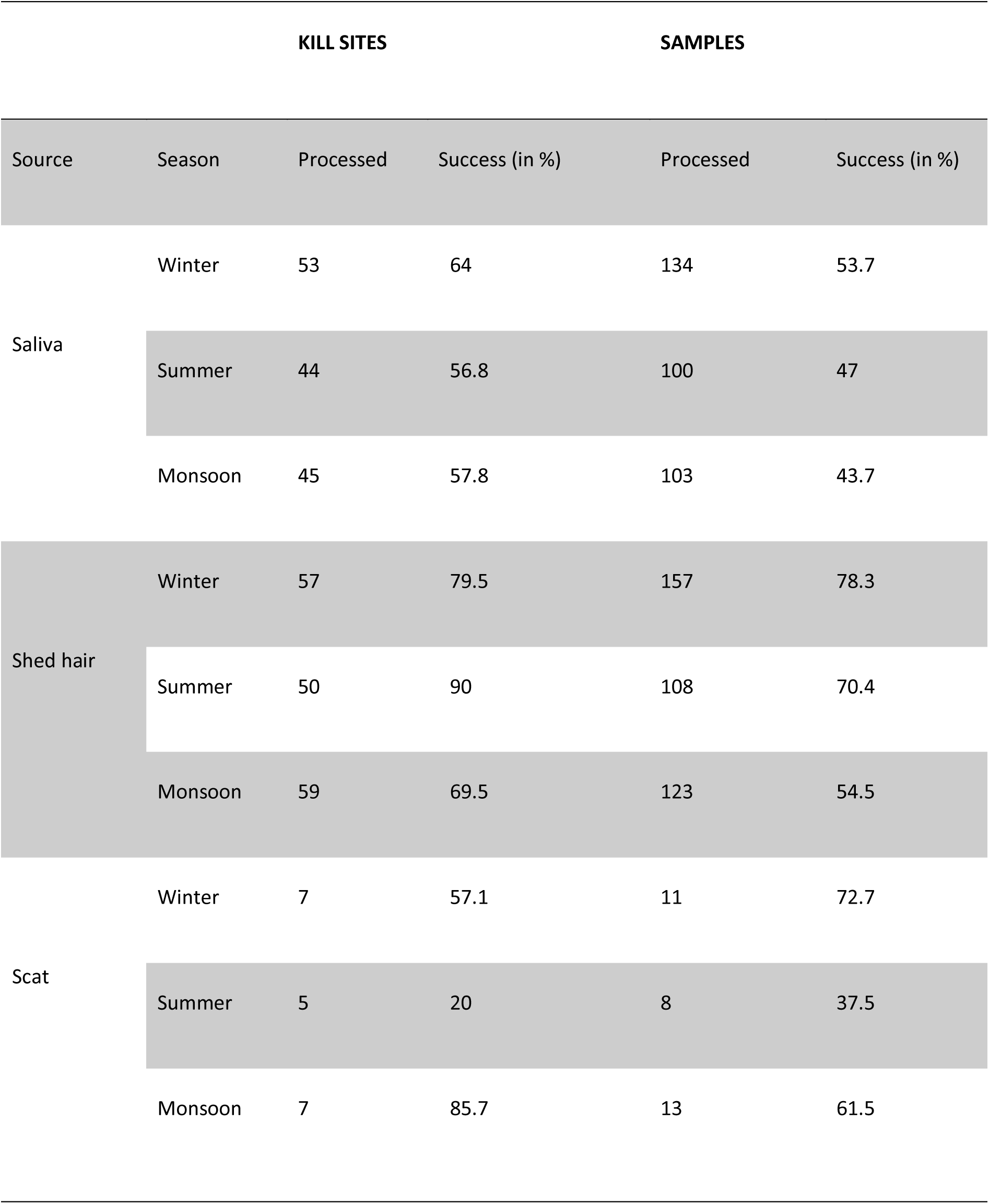
Summary of individual identification success for various sample types (saliva, shed hair and scat) across seasons (winter, summer and monsoon).

**Table 4:**
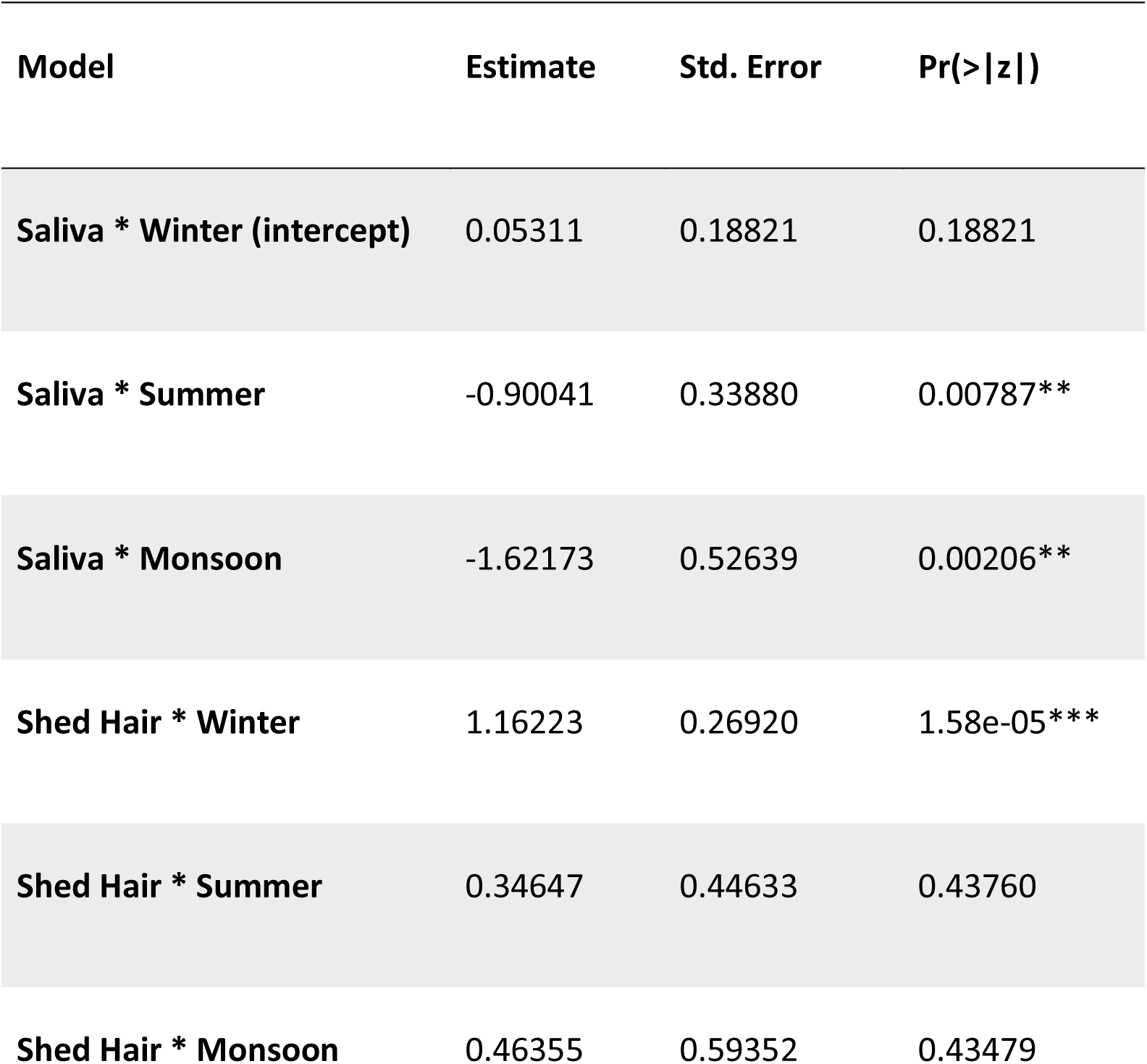
Generalised Linear Models (GLM) results examining the influence of sample type (saliva and shed hair) and season (winter, summer and monsoon) on individual identification success.

| Model | Estimate | Std. Error | Pr(>\|z\|) |
| --- | --- | --- | --- |
| Saliva * Winter (intercept) | 0.05311 | 0.18821 | 0.18821 |
| Saliva * Summer | -0.90041 | 0.33880 | 0.00787** |
| Saliva * Monsoon | -1.62173 | 0.52639 | 0.00206** |
| Shed Hair * Winter | 1.16223 | 0.26920 | 1.58e-05*** |
| Shed Hair * Summer | 0.34647 | 0.44633 | 0.43760 |
| Shed Hair * Monsoon | 0.46355 | 0.59352 | 0.43479 |

Based on the estimated mean confidence interval, we infer that the number of saliva samples required for confident individual identification from a kill site (with 90% probability) is 16, six and four samples in monsoon, summer, and winter, respectively (Supplementary Table S2). For shed hair samples, the numbers are lower overall, with five, three and two samples required in the monsoon, summer and winter, respectively.

### 3.4 Preliminary insights on individual engagement in depredation

While samples from 169 kill sites had more than 40 SNPs, individual tigers were identified at 155 of the 198 kill sites where tigers were detected. Eighty-seven unique individuals were identified from these sites. Multiple individuals were detected at 19 sites. Forty-six per cent of the individuals were detected in only one kill site, while almost a third (29%) were at two kill sites. In other words, 75% of all tigers found at kill sites were at only one or two kill sites. Twenty individuals (23%) were detected at three to five kill sites, and only two individuals (2%) were detected at seven kills. Our sampling of kill sites was more intensive in Bandhavgarh, and we attempted to investigate spatial distributions of individual predators here (Figure 3).

**Figure 3:**
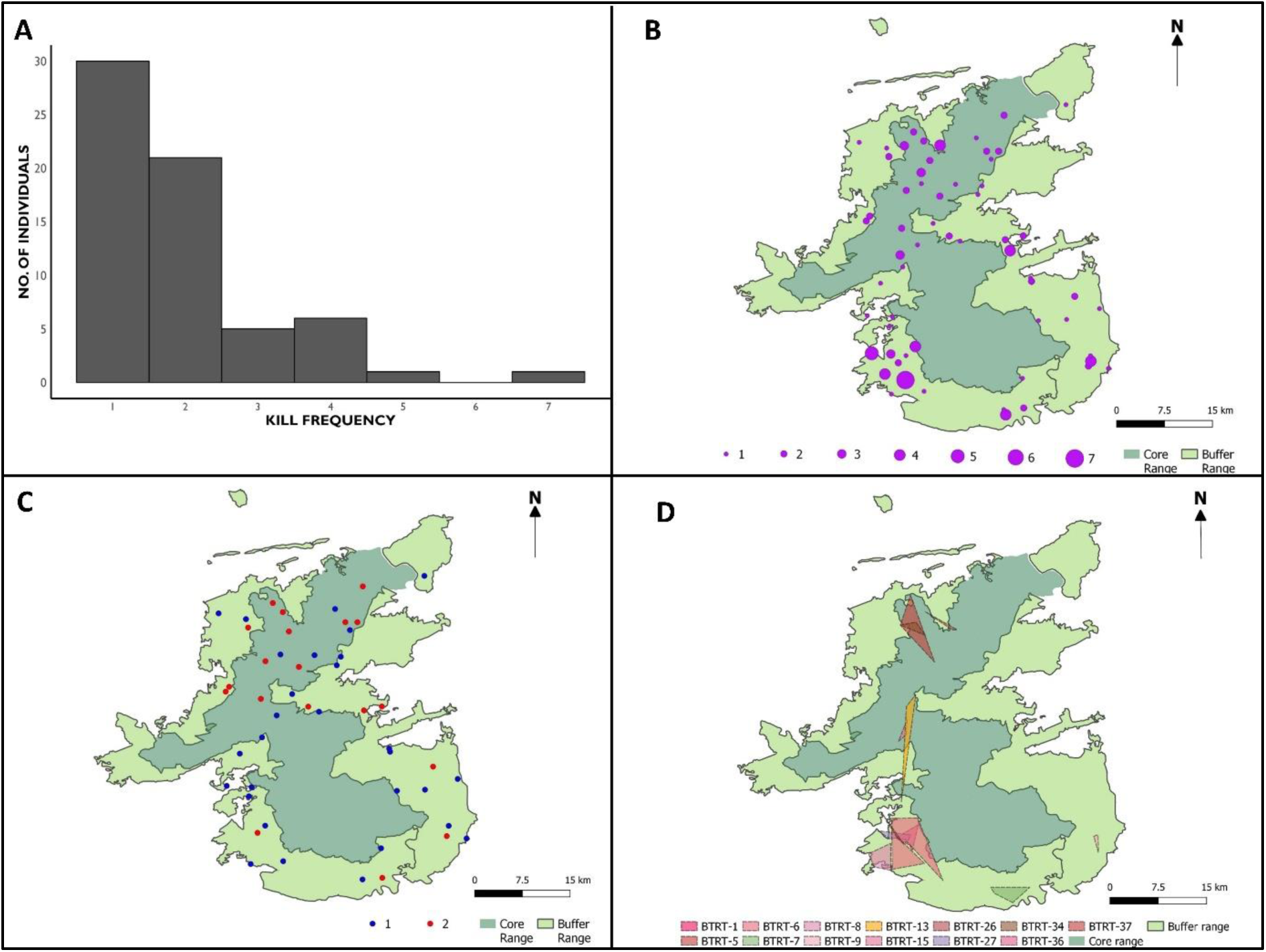
Summary of individuals identified across kills in Bandhavgarh Tiger Reserve. A) Histogram of individuals found across kill sites indicating majority of individuals were identified at one or two sites, B) frequency of individual occurrence across all kills, C) individuals identified (N = 51) at one or two kills D) occurrence polygon of individuals (N = 13) identified at 3 or more kills.

### 3.5 Predator assignment

We compiled predator assignments for both Bandhavgarh and Kanha. Tigers were detected at all 198 kill sites. By our defined categories, we were able to identify the ‘true predator’ at 151 kills, the ‘circumstantial predator’ and ‘ predator uncertain’ at the species level for 34 and 12 sites, respectively. Individual tigers were assigned as ‘true predators’ for 72 kills and as ‘circumstantial predators’ at 34 sites. Individuals were identified using multiple predation samples at 11 sites. All samples identified a unique individual in ten of these 11 sites. Two unique individuals were detected at one site using different predation samples, including saliva and shed hair. One of these individuals was detected at three other sites, while the second wasn’t recaptured at any other site. At 49 kill sites, individuals were identified as predator uncertain. Within these sites, individuals were classified as predator uncertain when identification was obtained from only one sample (38 kills) or when multiple individuals were detected (11 kills). Moreover, using the predation sample we successfully differentiated between predator and predator uncertain individuals at seven sites where multiple individuals were detected.

## 4. Discussion

Reliable identification of predating individuals is fundamental to understanding predation ecology, as well as in the management of human-wildlife conflict. Our study assessed the utility of trace DNA samples in predator identification while examining factors that influence success. We successfully identified tigers at all sampled sites, achieving a 100% success rate in species identification, and identified individual tigers at 86% of all kill sites. We find that the sample source and the sampling season are determinants of SNP typing and individual identification success. We develop a framework to assign confidence in predator assignments based on sample sources for individual identification. At 46% of kills, we identified true predator (high confidence), circumstantial predator (medium confidence) at 22% of sites, and predator uncertain (low confidence) at 32% of sites at individual level. We conclude that trace genetic samples, combined with systematic kill investigation, can successfully identify predators.

### 4.1 Season and sample source determine identification success

Shed hair samples consistently typed more SNPs and had higher individual identification success than saliva and scat samples, regardless of the season. Winter was observed to be the most favourable season for sampling. As predicted, the individual identification success rate per sample was lower during the monsoon season. We could not test the independent impact of rainfall and sampling time lag on success as rainfall was strongly associated with season, and we had a low sample size for delayed sampling events (> three days). While earlier studies in controlled settings have shown the negative impact of rainfall and time lag on DNA quality (Harms et al., 2015; Piaggio et al., 2020; Reddy et al., 2012), testing independent effects in field studies becomes challenging due to deposition of fresh saliva across feeding bouts. Despite declining individual identification success per sample, the success per kill remained consistent across seasons. This implies that collecting multiple samples from various sources at a single kill could help alleviate the seasonal impact. We therefore recommend the collection of more samples in monsoon followed by summer and winter. We endorse previous suggestions to collect samples promptly to prevent degradation and minimise the influence of scavengers (Ganz et al., 2023; Mumma et al., 2014). Additionally, we strongly recommend collecting multiple sources of samples, including predation samples (such as saliva from fatal wounds and shed hair from the point of kill), at all sites, as these samples aid in differentiation between predators and scavengers, ultimately strengthening confidence assignment in identification. We recognise that processing more samples can increase costs and recommend prioritising sample processing, such as predation samples, over others to reduce costs.

### 4.2 High proportion of individuals engage in depredation

Human-wildlife conflict poses a significant threat to conservation initiatives, and targeted removal of individuals is widely used as a strategy for conflict management. This approach, however, assumes unequal contributions of specific “problem” individuals in conflict engagement, essentially drawing from intra-species variations (Swan et al., 2017). We identified 87 individual tigers from 155 sites in Kanha and Bandhavgarh that were present at livestock kills. Considering the recent findings from the All India Tiger Estimation (Qureshi et al., 2023), which estimated Bandhavgarh’s tiger population at 165 individuals, our study identified ∼40% of these individuals at livestock kills. 75% of the individuals identified, in both Kanha and Bandhavgarh, were involved in less than three kills while accounting for 58% of kills. Establishing presence of problem individuals would require information on intra-population variation in diet and livestock depredation frequencies by individual tigers, resulting economic losses and the determinants of this variation. We acknowledge the limitations of our study to comment on the existence of “problem” individuals, and further research is required to validate their presence, especially in systems with large numbers of livestock kills such as ours. Moreover, we propose differential assessments of cases involving livestock depredation and human attacks when investigating potential problem individuals. Removal of individuals has consequences on population dynamics and raises ethical considerations. Therefore, for such interventions to be effective, it is crucial to establish the existence of problem individuals in the system and accurately identify the predator before resorting to targeted removal. Otherwise, targeted removal will be a tool for balancing conservation and political goals without serving its intended purpose of conflict reduction.

### 4.3 Conclusive identification of predator will require an adaptive and systematic sampling design

Genetic methods are more effective in predator identification than field-based methods (Ganz et al., 2023; Mumma et al., 2014). Samples can also be collected from human predation sites where use of conventional methods becomes a challenge and has ethical considerations (Pandey & Sharma, 2016). They are advantageous in identifying the individual predator even when visually unique individual markings are lacking. Trace DNA deposited during a predation event (such as saliva from predation wounds and shed hair at kill locations), essentially serves as evidence, whereas camera traps are typically deployed after the event. However, genetic and conventional methods are prone to misidentification, especially when multiple potential predators and scavengers are at the kill site (Ganz et al., 2023; Steffens et al., 2012; Verzuh et al., 2018). It is, therefore, crucial to triangulate information from various sources, including field investigations, camera trapping, genetic data, and data from collared individuals to identify predators reliably. Once a predation event is confirmed (see Cristescu et al., 2022), the kill site should be systematically explored and documented, followed by meticulous sample collection. To enhance the reliability of assessments, we recommend assigning confidence levels based on the information source and associated errors. We provide a framework for confidence assignment using genetic data by categorising samples according to their source and collection site. The highest confidence is attributed to samples likely deposited during a predation event, such as saliva samples from fatal wounds (indicated by haemorrhage) and shed hair samples from the inferred kill location. This framework can be adapted and modified in future studies to encompass various other sources of information like camera trap images from kill sites, prior knowledge of individual territories and radio telemetry data. We align with Cristescu et al. (2022) in recognising that reporting such evidence and subsequent assessments will enhance transparency and foster public support in decision-making.

### 4.4 Integration of molecular approaches into management

The broader acceptance and utilisation of genetic tools face challenges due to cold storage, extended processing times, and high sample processing costs (Khan & Tyagi, 2021). However, lysis buffers allow for sample storage at room temperature, addressing the cold storage requirement. Advancements in next-generation sequencing methods can further diminish processing time and costs (see Natesh et al., 2019). These methods also enable the real-time generation of high-quality data using non-invasive samples (Urban et al., 2023). We propose establishing a reference genetic database, particularly for individuals involved in conflicts. Furthermore, acknowledging that genetic information does not align with the visual identification typically required for management actions, this database can be linked to the physical identification of individuals. This can be achieved by simultaneously sampling kills for trace DNA and deploying camera traps at kill sites in addition to opportunistic sampling during animal sightings and the capture of individuals. This comprehensive database of individuals will enable a better understanding of predation ecology through robust individual identification and promote evidence-based conflict management.

## Acknowledgements

We acknowledge the role and support of the Madhya Pradesh Forest Department especially that of Kanha and Bandhavgarh Tiger Reserves. We thank Meghana Natesh, Vinay Sagar for their support during NGS data analysis and Divya Verma, Adeeba for helping with lab work. We thank Prasenjeet Yadav and Ankit Kacchwaha for their support during the fieldwork. Ashwin Vishwanathan and Ramya Roopa provided valuable inputs on statistical analysis of the data. We also would like to thank Jessica Luis for help with the amazing sketch of our field sampling scheme.

This research was supported by a National Geographic explorer grant to UR, HC and AH (NGS-62344R-19) and the NCBS/TIFR plan fund awarded to UR (Project Identification RTI 4006, Department of Atomic Energy, Government of India). The NCBS data cluster used is supported under project 12-R&D-TFR-5.04-0900, Department of Atomic Energy, Government of India. Field research permits were provided by the Madhya Pradesh Forest Department (No./Tech-1/2048 & No./Tech-1/7661). Biosafety approvals were obtained from the Institutional Biosafety Committee at NCBS (TIFR:NCBS:29-IBSC/UR1).

## Author Contributions

HC, AH, and UR conceived the ideas and designed methodology; HC, RD, AY and KP conducted field sampling; HC, DR and AP did laboratory work and NGS data analysis; SS and VR provided essential support for field sampling; HC wrote the first draft. All authors gave final approval for publication.

## Data availability

Raw sequence data and scripts for variant calling will be deposited at NCBI and Github, respectively, and will be made available upon acceptance.

## References

Armstrong, E. E., Khan, A., Taylor, R. W., Gouy, A., Greenbaum, G., Thiéry, A., Kang, J. T., Redondo, S. A., Prost, S., Barsh, G., Kaelin, C., Phalke, S., Chugani, A., Gilbert, M., Miquelle, D., Zachariah, A., Borthakur, U., Reddy, A., Louis, E., … Ramakrishnan, U. (2021). Recent Evolutionary History of Tigers Highlights Contrasting Roles of Genetic Drift and Selection. Molecular Biology and Evolution, 38(6), 2366–2379. 10.1093/molbev/msab032

Athreya, V., Odden, M., Linnell, J. D. C., & Karanth, K. U. (2011). Translocation as a Tool for Mitigating Conflict with Leopards in Human-Dominated Landscapes of India: Human-Leopard Conflicts. Conservation Biology, 25(1), 133–141. 10.1111/j.1523-1739.2010.01599.x

Awasthi, N., Kumar, U., Qureshi, Q., Pradhan, A., Chauhan, J. S., & Jhala, Y. V. (2016). Effect of human use, season and habitat on ungulate density in Kanha Tiger Reserve, Madhya Pradesh, India. Regional Environmental Change, 16(S1), 31–41. 10.1007/s10113-016-0953-z

Balme, G. A., Miller, J. R. B., Pitman, R. T., & Hunter, L. T. B. (2017). Caching reduces kleptoparasitism in a solitary, large felid. Journal of Animal Ecology, 86(3), 634–644. 10.1111/1365-2656.12654

Berezowska-Cnota, T., Konopiński, M. K., Bartoń, K., Bautista, C., Revilla, E., Naves, J., Biedrzycka, A., Fedyń, H., Fernández, N., Jastrzębski, T., Pirga, B., Viota, M., Wojtas, Z., & Selva, N. (2023). Individuality matters in human–wildlife conflicts: Patterns and fraction of damage-making brown bears in the north-eastern Carpathians. Journal of Applied Ecology, 60(6), 1127–1138. 10.1111/1365-2664.14388

Bhagavatula, J., & Singh, L. (2006). Genotyping faecal samples of Bengal tiger Panthera tigris tigris for population estimation: A pilot study. BMC Genetics, 7(1), 48. 10.1186/1471-2156-7-48

Blejwas, K. M., Williams, C. L., Shin, G. T., McCullough, D. R., & Jaeger, M. M. (2006). Salivary DNA Evidence Convicts Breeding Male Coyotes of Killing Sheep. The Journal of Wildlife Management, 70(4), 1087–1093.

Bolnick, D. I., Svanbäck, R., Fordyce, J. A., Yang, L. H., Davis, J. M., Hulsey, C. D., & Forister, M. L. (2003). The Ecology of Individuals: Incidence and Implications of Individual Specialization. The American Naturalist, 161(1), 1–28. 10.1086/343878

Caniglia, R., Mastrogiuseppe, L., Randi, E., & Fabbri, E. (2012). Caniglia et al. Wolf saliva FSIgen 2012 [dataset].

Champion, S. H. G., & Seth, S. K. (1968). A Revised Survey of the Forest Types of India. Manager of Publications.

Chundawat, R. S., Sharma, K., Gogate, N., Malik, P. K., & Vanak, A. T. (2016). Size matters: Scale mismatch between space use patterns of tigers and protected area size in a Tropical Dry Forest. Biological Conservation, 197, 146–153. 10.1016/j.biocon.2016.03.004

Cristescu, B., Elbroch, L. M., Forrester, T. D., Allen, M. L., Spitz, D. B., Wilmers, C. C., & Wittmer, H. U. (2022). Standardizing protocols for determining the cause of mortality in wildlife studies. Ecology and Evolution, 12(6), e9034. 10.1002/ece3.9034

DePristo, M. A., Banks, E., Poplin, R., Garimella, K. V., Maguire, J. R., Hartl, C., Philippakis, A. A., del Angel, G., Rivas, M. A., Hanna, M., McKenna, A., Fennell, T. J., Kernytsky, A. M., Sivachenko, A. Y., Cibulskis, K., Gabriel, S. B., Altshuler, D., & Daly, M. J. (2011). A framework for variation discovery and genotyping using next-generation DNA sequencing data. Nature Genetics, 43(5), Article 5. 10.1038/ng.806

Dickman, C. R., & Newsome, T. M. (2015). Individual hunting behaviour and prey specialisation in the house cat Felis catus: Implications for conservation and management. Applied Animal Behaviour Science, 173, 76–87. 10.1016/j.applanim.2014.09.021

Estes, J. A., Riedman, M. L., Staedler, M. M., Tinker, M. T., & Lyon, B. E. (2003). Individual variation in prey selection by sea otters: Patterns, causes and implications. Journal of Animal Ecology, 72(1), 144–155. 10.1046/j.1365-2656.2003.00690.x

Fotedar, S., Lukehurst, S., Jackson, G., & Snow, M. (2019). Molecular tools for identification of shark species involved in depredation incidents in Western Australian fisheries. PLOS ONE, 14(1), e0210500. 10.1371/journal.pone.0210500

Ganz, T. R., DeVivo, M. T., Reese, E. M., & Prugh, L. R. (2023). Wildlife whodunnit: Forensic identification of predators to inform wildlife management and conservation. Wildlife Society Bulletin, 47(1), e1386. 10.1002/wsb.1386

Goodrich, J. M. (2010). Human–tiger conflict: A review and call for comprehensive plans. Integrative Zoology, 5(4), 300–312. 10.1111/j.1749-4877.2010.00218.x

Harms, V., Nowak, C., Carl, S., & Muñoz-Fuentes, V. (2015). Experimental evaluation of genetic predator identification from saliva traces on wildlife kills. Journal of Mammalogy, 96(1), 138–143. 10.1093/jmammal/gyu014

Karanth, K. U., & Nichols, J. D. (Eds.). (2002). Monitoring tigers and their prey: A manual for wildlife researchers, managers and conservationists in tropical Asia. Centre for Wildlife Studies; USGS Publications Warehouse. https://pubs.usgs.gov/publication/5200258

Khan, A., & Tyagi, A. (2021). Considerations for Initiating a Wildlife Genomics Research Project in South and South-East Asia. Journal of the Indian Institute of Science, 101(2), 243–256. 10.1007/s41745-021-00243-3

Li, H. (2011). A statistical framework for SNP calling, mutation discovery, association mapping and population genetical parameter estimation from sequencing data. Bioinformatics, 27(21), 2987–2993. 10.1093/bioinformatics/btr509

Li, H. (2013). Aligning sequence reads, clone sequences and assembly contigs with BWA-MEM (arXiv:1303.3997). arXiv. http://arxiv.org/abs/1303.3997

Linnell, J. D. C., Odden, J., Smith, M. E., Aanes, R., & Swenson, J. E. (1999). Large Carnivores That Kill Livestock: Do “Problem Individuals” Really Exist? Wildlife Society Bulletin (1973-2006), 27(3), 698–705.

Longmire, J. L., Maltbie, M., Baker, R. J., & Texas Tech University. (1997). Use of “Lysis Buffer” in DNA isolation and its implication for museum collections. 10.5962/bhl.title.143318

Mumma, M. A., Soulliere, C. E., Mahoney, S. P., & Waits, L. P. (2014). Enhanced understanding of predator-prey relationships using molecular methods to identify predator species, individual and sex. Molecular Ecology Resources, 14(1), 100–108. 10.1111/1755-0998.12153

Nakamura, M., Godinho, R., Rio-Maior, H., Roque, S., Kaliontzopoulou, A., Bernardo, J., Castro, D., Lopes, S., Petrucci-Fonseca, F., & Álvares, F. (2017). Evaluating the predictive power of field variables for species and individual molecular identification on wolf noninvasive samples. European Journal of Wildlife Research, 63(3), 53. 10.1007/s10344-017-1112-7

Natesh, M., Taylor, R. W., Truelove, N. K., Hadly, E. A., Palumbi, S. R., Petrov, D. A., & Ramakrishnan, U. (2019). Empowering conservation practice with efficient and economical genotyping from poor quality samples. Methods in Ecology and Evolution, 10(6), 853–859. 10.1111/2041-210X.13173

Negi, H. S., & Shukla, R. (2011). Tiger Conservation Plan for Kanha tiger reserve. Office of the Field Director Kanha Tiger Reserve Mandla Madhya Pradesh.

Nichols, R. V., Königsson, H., Danell, K., & Spong, G. (2012). Browsed twig environmental DNA: Diagnostic PCR to identify ungulate species. Molecular Ecology Resources, 12(6), 983– 989. 10.1111/j.1755-0998.2012.03172.x

Pandey, P., & Sharma, V. (2016). Curtailing Human-Leopard Conflict Using Wildlife Forensics: A Case Study from Himachal Pradesh, India. Journal of Forensic Research, 7(3). 10.4172/2157-7145.1000331

Périquet, S., Fritz, H., & Revilla, E. (2015). The Lion King and the Hyaena Queen: Large carnivore interactions and coexistence. Biological Reviews, 90(4), 1197–1214. 10.1111/brv.12152

Piaggio, A. J., Shriner, S. A., Young, J. K., Griffin, D. L., Callahan, P., Wostenberg, D. J., Gese, E. M., & Hopken, M. W. (2020). DNA persistence in predator saliva from multiple species and methods for optimal recovery from depredated carcasses. Journal of Mammalogy, 101(1), 298–306. 10.1093/jmammal/gyz156

Purcell, S., Neale, B., Todd-Brown, K., Thomas, L., Ferreira, M. A. R., Bender, D., Maller, J., Sklar, P., De Bakker, P. I. W., Daly, M. J., & Sham, P. C. (2007). PLINK: A Tool Set for Whole-Genome Association and Population-Based Linkage Analyses. The American Journal of Human Genetics, 81(3), 559–575. 10.1086/519795

Qamar Qureshi, Yadvendradev V. Jhala, Satya P. Yadav and Amit Mallick (eds) 2023. Status of tigers, co-predators and prey in India, 2022. National Tiger Conservation Authority, Government of India, New Delhi, and Wildlife Institute of India, Dehradun.

Reddy, P. A., Bhavanishankar, M., Bhagavatula, J., Harika, K., Mahla, R. S., & Shivaji, S. (2012). Improved Methods of Carnivore Faecal Sample Preservation, DNA Extraction and Quantification for Accurate Genotyping of Wild Tigers. PLOS ONE, 7(10), e46732. 10.1371/journal.pone.0046732

Sagar, V., Kaelin, C. B., Natesh, M., Reddy, P. A., Mohapatra, R. K., Chhattani, H., Thatte, P., Vaidyanathan, S., Biswas, S., Bhatt, S., Paul, S., Jhala, Y. V., Verma, M. M., Pandav, B., Mondol, S., Barsh, G. S., Swain, D., & Ramakrishnan, U. (2021). High frequency of an otherwise rare phenotype in a small and isolated tiger population. Proceedings of the National Academy of Sciences, 118(39), e2025273118. 10.1073/pnas.2025273118

Scholz, C., Firozpoor, J., Kramer-Schadt, S., Gras, P., Schulze, C., Kimmig, S. E., Voigt, C. C., & Ortmann, S. (2020). Individual dietary specialization in a generalist predator: A stable isotope analysis of urban and rural red foxes. Ecology and Evolution, 10(16), 8855–8870. 10.1002/ece3.6584

Sinha, N. (2018). To Kill a Tigress. 53(48). https://www.epw.in/journal/2018/48/commentary/kill-tigress.html

Steffens, K. E., Sanders, M. D., Gleenson, D. M., Pullen, K. M., & Stowe, C. J. (2012). Identification of predators at black-fronted tern Chlidonias albostriatus nests, using mtDNA analysis and digital video recorders. New Zealand Journal of Ecology, 36(1), 48–55.

Sundqvist, A.-K., Ellegren, H., & Vilà, C. (2008). Wolf or dog? Genetic identification of predators from saliva collected around bite wounds on prey. Conservation Genetics, 9(5), 1275–1279. 10.1007/s10592-007-9454-4

Swan, G. J. F., Redpath, S. M., Bearhop, S., & McDonald, R. A. (2017). Ecology of Problem Individuals and the Efficacy of Selective Wildlife Management. Trends in Ecology & Evolution, 32(7), 518–530. 10.1016/j.tree.2017.03.011

Treves, A., & Karanth, K. U. (2003). Human-Carnivore Conflict and Perspectives on Carnivore Management Worldwide. Conservation Biology, 17(6), 1491–1499. 10.1111/j.1523-1739.2003.00059.x

Urban, L., Miller, A. K., Eason, D., Vercoe, D., Shaffer, M., Wilkinson, S. P., Jeunen, G.-J., Gemmell, N. J., & Digby, A. (2023). Non-invasive real-time genomic monitoring of the critically endangered kākāpō. eLife, 12. 10.7554/eLife.84553.1

Verzuh, T., Bergman, D. L., Bender, S. C., Dwire, M., & Breck, S. W. (2018). Intercanine width measurements to aid predation investigations: A comparison between sympatric native and non-native carnivores in the Mexican wolf recovery area. Journal of Mammalogy, 99(6), 1405–1410. 10.1093/jmammal/gyy145

Voigt, C. C., Krofel, M., Menges, V., Wachter, B., & Melzheimer, J. (2018). Sex-specific dietary specialization in a terrestrial apex predator, the leopard, revealed by stable isotope analysis. Journal of Zoology, 306(1), 1–7. 10.1111/jzo.12566

Wheat, R. E., Allen, J. M., Miller, S. D. L., Wilmers, C. C., & Levi, T. (2016). Environmental DNA from Residual Saliva for Efficient Noninvasive Genetic Monitoring of Brown Bears (Ursus arctos). PLOS ONE, 11(11), e0165259. 10.1371/journal.pone.0165259

Woodroffe, R., & Frank, L. G. (2005). Lethal control of African lions (Panthera leo): Local and regional population impacts. Animal Conservation, 8(1), 91–98. 10.1017/S1367943004001829

