## Supplementary Figure S1, Supplementary Table S1, Supplementary Figure S2, Supplementary Figure S3, Supplementary Table S2 for "Trace DNA from kill sites identifies predating tigers"

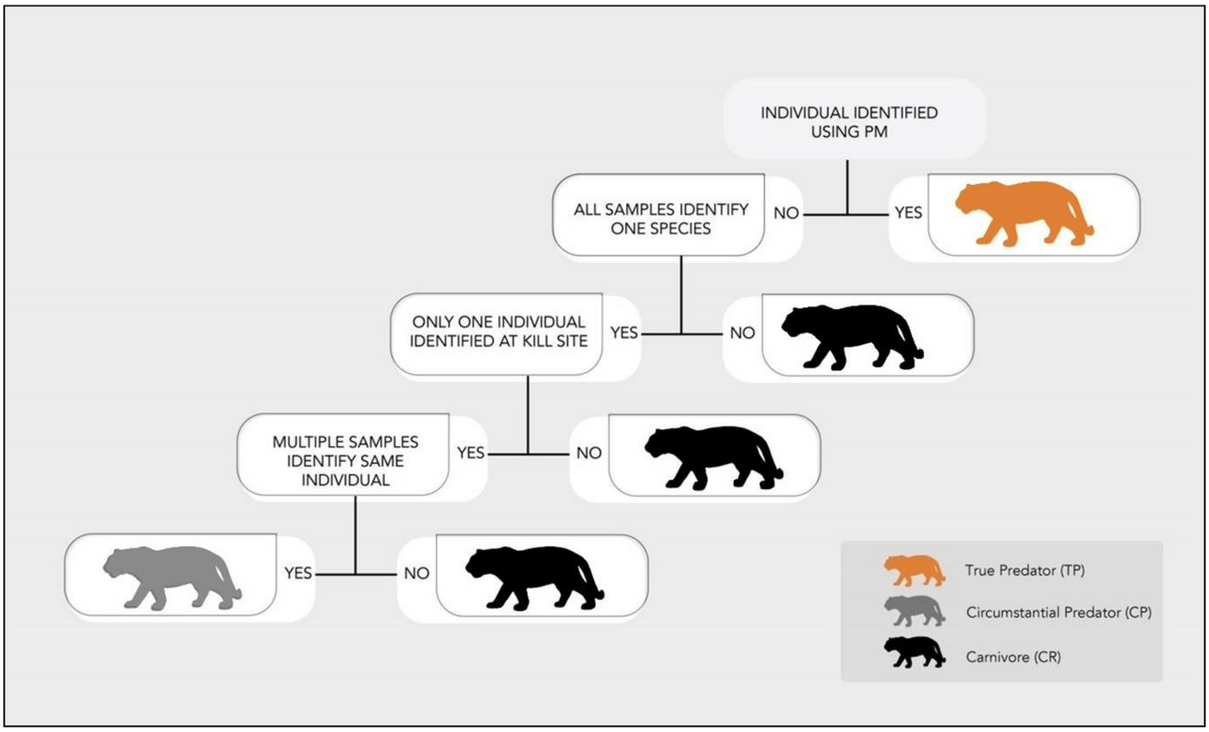


**Figure S1:** Schematic flow chart for categorisation of individual tigers.

**Table S1:** Table summarising number of samples collected from individual kill sites in Kanha and Bandhavgarh.

| Kill ID | Season | Reserve | Number of samples collected | | | |
|---|---|---|---|---|---|---|
| | | | Saliva | Scat | Shed hair | Total |
| C01-KTR17-118 | Monsoon | Kanha | 2 | | 1 | 3 |
| C01-KTR17-119 | Monsoon | Kanha | | | 1 | 1 |
| C01-KTR17-120 | Monsoon | Kanha | 2 | | 1 | 3 |
| C01-KTR17-121 | Monsoon | Kanha | 3 | 2 | | 5 |
| C01-KTR17-122 | Monsoon | Kanha | 3 | 1 | 1 | 5 |
| C01-KTR17-123 | Monsoon | Kanha | 3 | | | 3 |
| C01-KTR17-125 | Monsoon | Kanha | 3 | | 1 | 4 |
| C01-KTR17-126 | Monsoon | Kanha | 3 | | 1 | 4 |
| C01-KTR17-127 | Monsoon | Kanha | | | 1 | 1 |
| C01-KTR17-128 | Monsoon | Kanha | 3 | | | 3 |
| C01-KTR17-129 | Monsoon | Kanha | 3 | | 1 | 4 |
| C01-KTR17-131 | Monsoon | Kanha | 2 | | | 2 |
| C01-KTR17-132 | Monsoon | Kanha | 3 | | 1 | 4 |
| C01-KTR17-136 | Monsoon | Kanha | 2 | | | 2 |
| C01-KTR17-137 | Monsoon | Kanha | 2 | | | 2 |

| | | | | | | |
|---|---|---|---|---|---|---|
| C01-KTR17-138 | Monsoon | Kanha | 3 | | | 3 |
| C01-KTR17-139 | Monsoon | Kanha | 3 | | | 3 |
| C01-KTR17-140 | Monsoon | Kanha | 3 | | | 3 |
| C01-KTR17-142 | Monsoon | Kanha | 3 | | | 3 |
| C01-KTR17-143 | Monsoon | Kanha | 3 | | | 3 |
| C01-KTR17-144 | Monsoon | Kanha | 2 | 2 | | 4 |
| C01-KTR17-145 | Monsoon | Kanha | 3 | | | 3 |
| C01-KTR17-146 | Monsoon | Kanha | | | 1 | 1 |
| C01-KTR17-153 | Monsoon | Kanha | 3 | | | 3 |
| C01-KTR17-157 | Monsoon | Kanha | 3 | | | 3 |
| C01-KTR17-159 | Monsoon | Kanha | 3 | | 1 | 4 |
| C01-KTR17-162 | Monsoon | Kanha | 1 | | 1 | 2 |
| C01-KTR17-163 | Monsoon | Kanha | 1 | | | 1 |
| C01-KTR17-166 | Monsoon | Kanha | 3 | | 2 | 5 |
| C01-KTR17-168 | Monsoon | Kanha | 3 | | 1 | 4 |
| C01-KTR17-174 | Monsoon | Kanha | 3 | | | 3 |
| C02-BTR21-154 | Monsoon | Bandhavgarh | 2 | | 3 | 5 |
| C02-BTR21-157 | Monsoon | Bandhavgarh | 1 | | 2 | 3 |
| C02-BTR21-162 | Monsoon | Bandhavgarh | | | 1 | 1 |

| | | | | | | |
|---|---|---|---|---|---|---|
| C02-BTR21-164 | Monsoon | Bandhavgarh | | | 1 | 1 |
| C02-BTR21-170 | Monsoon | Bandhavgarh | | | 1 | 1 |
| C02-BTR21-172 | Monsoon | Bandhavgarh | | | 1 | 1 |
| C02-BTR21-175 | Monsoon | Bandhavgarh | | | 2 | 2 |
| C02-BTR21-179 | Monsoon | Bandhavgarh | | | 1 | 1 |
| C02-BTR21-180 | Monsoon | Bandhavgarh | | | 3 | 3 |
| C02-BTR21-184 | Monsoon | Bandhavgarh | | | 3 | 3 |
| C02-BTR21-191 | Monsoon | Bandhavgarh | | | 1 | 1 |
| C02-BTR21-200 | Monsoon | Bandhavgarh | 3 | | 3 | 6 |
| C02-BTR21-203 | Monsoon | Bandhavgarh | 1 | | 1 | 2 |
| C02-BTR21-209 | Monsoon | Bandhavgarh | 3 | | 4 | 7 |
| C02-BTR21-211 | Monsoon | Bandhavgarh | | | 3 | 3 |
| C02-BTR21-218 | Monsoon | Bandhavgarh | | | 3 | 3 |
| C02-BTR21-224 | Monsoon | Bandhavgarh | | | 3 | 3 |
| C02-BTR21-227 | Monsoon | Bandhavgarh | 1 | | 2 | 3 |
| C02-BTR21-228 | Monsoon | Bandhavgarh | 1 | 2 | 3 | 6 |
| C02-BTR21-234 | Monsoon | Bandhavgarh | | | 3 | 3 |
| C02-BTR21-236 | Monsoon | Bandhavgarh | | | 3 | 3 |
| C02-BTR21-237 | Monsoon | Bandhavgarh | 2 | | 3 | 5 |

| | | | | | | |
|---|---|---|---|---|---|---|
| C04-BTR21-050 | Monsoon | Bandhavgarh | | | 2 | 2 |
| C04-BTR21-051 | Monsoon | Bandhavgarh | | | 3 | 3 |
| C04-BTR21-053 | Monsoon | Bandhavgarh | | | 4 | 4 |
| C04-BTR21-054 | Monsoon | Bandhavgarh | | | 2 | 2 |
| C04-BTR21-066 | Monsoon | Bandhavgarh | 1 | | 2 | 3 |
| C04-BTR21-074 | Monsoon | Bandhavgarh | 1 | | 1 | 2 |
| C04-BTR21-075 | Monsoon | Bandhavgarh | | | 5 | 5 |
| C04-BTR21-076 | Monsoon | Bandhavgarh | | | 1 | 1 |
| C04-BTR21-077 | Monsoon | Bandhavgarh | | | 4 | 4 |
| C04-BTR21-078 | Monsoon | Bandhavgarh | 3 | | 2 | 5 |
| C04-BTR21-079 | Monsoon | Bandhavgarh | | | 2 | 2 |
| C04-BTR21-083 | Monsoon | Bandhavgarh | 1 | | | 1 |
| C04-BTR21-085 | Monsoon | Bandhavgarh | | | 2 | 2 |
| C04-BTR21-086 | Monsoon | Bandhavgarh | | | 1 | 1 |
| C04-BTR21-087 | Monsoon | Bandhavgarh | 1 | 2 | 2 | 5 |
| C04-BTR21-088 | Monsoon | Bandhavgarh | 2 | 3 | 1 | 6 |
| C04-BTR21-089 | Monsoon | Bandhavgarh | 2 | 1 | 3 | 6 |
| C04-BTR21-090 | Monsoon | Bandhavgarh | 2 | | 1 | 3 |
| C04-BTR21-091 | Monsoon | Bandhavgarh | | | 4 | 4 |

| | | | | | | |
|---|---|---|---|---|---|---|
| C04-BTR21-092 | Monsoon | Bandhavgarh | 2 | | 2 | 4 |
| C04-BTR21-094 | Monsoon | Bandhavgarh | | | 4 | 4 |
| C04-BTR21-095 | Monsoon | Bandhavgarh | | | 3 | 3 |
| C04-BTR21-097 | Monsoon | Bandhavgarh | | | 4 | 4 |
| C04-BTR21-102 | Monsoon | Bandhavgarh | | | 3 | 3 |
| C04-BTR21-99 | Monsoon | Bandhavgarh | | | 4 | 4 |
| C01-KTR17-19 | Summer | Kanha | 5 | | 1 | 6 |
| C01-KTR17-20 | Summer | Kanha | 4 | | | 4 |
| C01-KTR17-32 | Summer | Kanha | 2 | | | 2 |
| C01-KTR17-33 | Summer | Kanha | 3 | 1 | | 4 |
| C01-KTR17-34 | Summer | Kanha | 3 | 4 | 2 | 9 |
| C01-KTR17-35 | Summer | Kanha | | | 2 | 2 |
| C01-KTR17-36 | Summer | Kanha | | | 1 | 1 |
| C01-KTR17-38 | Summer | Kanha | 3 | | 2 | 5 |
| C01-KTR17-43 | Summer | Kanha | 1 | | | 1 |
| C01-KTR17-51 | Summer | Kanha | 3 | | | 3 |
| C01-KTR17-53 | Summer | Kanha | 3 | | | 3 |
| C01-KTR17-54 | Summer | Kanha | | | 2 | 2 |
| C01-KTR17-55 | Summer | Kanha | 3 | | 3 | 6 |

| | | | | | | |
|---|---|---|---|---|---|---|
| C01-KTR17-57 | Summer | Kanha | 3 | | | 3 |
| C01-KTR17-58 | Summer | Kanha | 3 | | 1 | 4 |
| C01-KTR17-97 | Summer | Kanha | 2 | | | 2 |
| C01-KTR17-98 | Summer | Kanha | 1 | | | 1 |
| C01-KTR17-99 | Summer | Kanha | 1 | | | 1 |
| C02-BTR21-100 | Summer | Bandhavgarh | 2 | | 1 | 3 |
| C02-BTR21-102 | Summer | Bandhavgarh | 1 | | 2 | 3 |
| C02-BTR21-105 | Summer | Bandhavgarh | 1 | 1 | 2 | 4 |
| C02-BTR21-112 | Summer | Bandhavgarh | | | 4 | 4 |
| C02-BTR21-113 | Summer | Bandhavgarh | | | 3 | 3 |
| C02-BTR21-118 | Summer | Bandhavgarh | 2 | | 3 | 5 |
| C02-BTR21-127 | Summer | Bandhavgarh | 2 | | 1 | 3 |
| C02-BTR21-142 | Summer | Bandhavgarh | 2 | | 1 | 3 |
| C02-BTR21-143 | Summer | Bandhavgarh | 3 | | 2 | 5 |
| C02-BTR21-146 | Summer | Bandhavgarh | 3 | | 1 | 4 |
| C02-BTR21-37 | Summer | Bandhavgarh | | | 1 | 1 |
| C02-BTR21-43 | Summer | Bandhavgarh | 3 | | 1 | 4 |
| C02-BTR21-45 | Summer | Bandhavgarh | 2 | | 2 | 4 |
| C02-BTR21-46 | Summer | Bandhavgarh | 3 | | 1 | 4 |

| | | | | | | |
|---|---|---|---|---|---|---|
| C02-BTR21-50 | Summer | Bandhavgarh | 2 | | 2 | 4 |
| C02-BTR21-54 | Summer | Bandhavgarh | 3 | | 2 | 5 |
| C02-BTR21-56 | Summer | Bandhavgarh | 2 | | 1 | 3 |
| C02-BTR21-57 | Summer | Bandhavgarh | 2 | | 1 | 3 |
| C02-BTR21-59 | Summer | Bandhavgarh | 2 | | 2 | 4 |
| C02-BTR21-61 | Summer | Bandhavgarh | 1 | | 2 | 3 |
| C02-BTR21-62 | Summer | Bandhavgarh | 2 | | 3 | 5 |
| C02-BTR21-64 | Summer | Bandhavgarh | 2 | | 2 | 4 |
| C02-BTR21-65 | Summer | Bandhavgarh | 2 | | 1 | 3 |
| C02-BTR21-74 | Summer | Bandhavgarh | | 2 | 3 | 5 |
| C02-BTR21-78 | Summer | Bandhavgarh | 1 | 1 | 2 | 4 |
| C02-BTR21-94 | Summer | Bandhavgarh | 2 | | 2 | 4 |
| C02-BTR21-98 | Summer | Bandhavgarh | 1 | | 2 | 3 |
| C04-BTR21-035 | Summer | Bandhavgarh | 2 | | 2 | 4 |
| C04-BTR21-038 | Summer | Bandhavgarh | | | 6 | 6 |
| C04-BTR21-040 | Summer | Bandhavgarh | 2 | | 3 | 5 |
| C04-BTR22-159 | Summer | Bandhavgarh | | | 3 | 3 |
| C04-BTR22-163 | Summer | Bandhavgarh | | | 3 | 3 |
| C04-BTR22-166 | Summer | Bandhavgarh | | | 4 | 4 |

| | | | | | | |
|---|---|---|---|---|---|---|
| C04-BTR22-167 | Summer | Bandhavgarh | | | 3 | 3 |
| C04-BTR22-168 | Summer | Bandhavgarh | | | 4 | 4 |
| C04-BTR22-169 | Summer | Bandhavgarh | | | 3 | 3 |
| C04-BTR22-175 | Summer | Bandhavgarh | | | 2 | 2 |
| C04-BTR22-177 | Summer | Bandhavgarh | | | 3 | 3 |
| C04-BTR22-178 | Summer | Bandhavgarh | 3 | | 3 | 6 |
| C04-BTR22-179 | Summer | Bandhavgarh | 3 | | 3 | 6 |
| C04-BTR22-181 | Summer | Bandhavgarh | 3 | | 2 | 5 |
| C04-BTR22-182 | Summer | Bandhavgarh | 1 | | 2 | 3 |
| C01-KTR17-03 | Winter | Kanha | 3 | | 1 | 4 |
| C01-KTR17-07 | Winter | Kanha | 3 | | | 3 |
| C01-KTR17-11 | Winter | Kanha | 3 | | 1 | 4 |
| C01-KTR17-12 | Winter | Kanha | 3 | | 1 | 4 |
| C01-KTR17-13 | Winter | Kanha | 3 | | | 3 |
| C01-KTR17-14 | Winter | Kanha | 3 | | 1 | 4 |
| C01-KTR17-15 | Winter | Kanha | 3 | 5 | 3 | 11 |
| C02-BTR21-242 | Winter | Bandhavgarh | 4 | | 2 | 6 |
| C02-BTR21-243 | Winter | Bandhavgarh | 3 | | 2 | 5 |
| C02-BTR21-249 | Winter | Bandhavgarh | 3 | | 1 | 4 |

| | | | | | | |
|---|---|---|---|---|---|---|
| C02-BTR21-252 | Winter | Bandhavgarh | 3 | | 3 | 6 |
| C02-BTR21-255 | Winter | Bandhavgarh | 2 | | 3 | 5 |
| C02-BTR21-257 | Winter | Bandhavgarh | | | 3 | 3 |
| C02-BTR21-258 | Winter | Bandhavgarh | 3 | | 1 | 4 |
| C02-BTR21-260 | Winter | Bandhavgarh | 2 | | 4 | 6 |
| C02-BTR21-263 | Winter | Bandhavgarh | 5 | | 3 | 8 |
| C02-BTR21-265 | Winter | Bandhavgarh | 4 | | 4 | 8 |
| C02-BTR21-271 | Winter | Bandhavgarh | 5 | | 3 | 8 |
| C02-BTR21-272 | Winter | Bandhavgarh | 3 | | 3 | 6 |
| C02-BTR21-277 | Winter | Bandhavgarh | 3 | | 4 | 7 |
| C02-BTR21-278 | Winter | Bandhavgarh | 2 | | 1 | 3 |
| C02-BTR21-283 | Winter | Bandhavgarh | 2 | | 4 | 6 |
| C02-BTR21-290 | Winter | Bandhavgarh | 5 | 1 | 7 | 13 |
| C02-BTR21-299 | Winter | Bandhavgarh | 1 | 1 | 1 | 3 |
| C02-BTR21-300 | Winter | Bandhavgarh | 2 | | 3 | 5 |
| C02-BTR21-303 | Winter | Bandhavgarh | | | 3 | 3 |
| C02-BTR21-304 | Winter | Bandhavgarh | | | 4 | 4 |
| C02-BTR21-308 | Winter | Bandhavgarh | 4 | 1 | 9 | 14 |
| C02-BTR21-309 | Winter | Bandhavgarh | 3 | | 2 | 5 |

| | | | | | | |
|---|---|---|---|---|---|---|
| C02-BTR21-315 | Winter | Bandhavgarh | 2 | 1 | 3 | 6 |
| C02-BTR21-317 | Winter | Bandhavgarh | 3 | | 4 | 7 |
| C02-BTR21-318 | Winter | Bandhavgarh | 3 | | 3 | 6 |
| C02-BTR21-322 | Winter | Bandhavgarh | 2 | | 4 | 6 |
| C02-BTR21-328 | Winter | Bandhavgarh | 2 | | 3 | 5 |
| C02-BTR21-337 | Winter | Bandhavgarh | 3 | | 3 | 6 |
| C02-BTR21-339 | Winter | Bandhavgarh | 3 | | 1 | 4 |
| C02-BTR21-340 | Winter | Bandhavgarh | 2 | | 3 | 5 |
| C04-BTR21-119 | Winter | Bandhavgarh | | | 3 | 3 |
| C04-BTR21-123 | Winter | Bandhavgarh | | | 4 | 4 |
| C04-BTR21-125 | Winter | Bandhavgarh | 2 | | 5 | 7 |
| C04-BTR21-126 | Winter | Bandhavgarh | | 1 | 3 | 4 |
| C04-BTR21-127 | Winter | Bandhavgarh | 2 | | 6 | 8 |
| C04-BTR21-129 | Winter | Bandhavgarh | 2 | | 1 | 3 |
| C04-BTR21-130 | Winter | Bandhavgarh | 2 | | 2 | 4 |
| C04-BTR21-131 | Winter | Bandhavgarh | 2 | | 1 | 3 |
| C04-BTR21-132 | Winter | Bandhavgarh | 3 | 1 | 3 | 7 |
| C04-BTR22-133 | Winter | Bandhavgarh | 2 | | 1 | 3 |
| C04-BTR22-134 | Winter | Bandhavgarh | 2 | | 1 | 3 |

| | | | | | | |
|---|---|---|---|---|---|---|
| C04-BTR22-135 | Winter | Bandhavgarh | 2 | | 2 | 4 |
| C04-BTR22-136 | Winter | Bandhavgarh | 2 | | 1 | 3 |
| C04-BTR22-137 | Winter | Bandhavgarh | 2 | | 2 | 4 |
| C04-BTR22-138 | Winter | Bandhavgarh | 2 | | 1 | 3 |
| C04-BTR22-139 | Winter | Bandhavgarh | 2 | | 3 | 5 |
| C04-BTR22-144 | Winter | Bandhavgarh | 2 | | 2 | 4 |
| C04-BTR22-145 | Winter | Bandhavgarh | 2 | | 4 | 6 |
| C04-BTR22-147 | Winter | Bandhavgarh | 2 | | 4 | 6 |
| C04-BTR22-148 | Winter | Bandhavgarh | 1 | | 2 | 3 |
| C04-BTR22-149 | Winter | Bandhavgarh | 2 | | 2 | 4 |
| C04-BTR22-151 | Winter | Bandhavgarh | 2 | | 2 | 4 |
| C04-BTR22-152 | Winter | Bandhavgarh | 1 | | 2 | 3 |

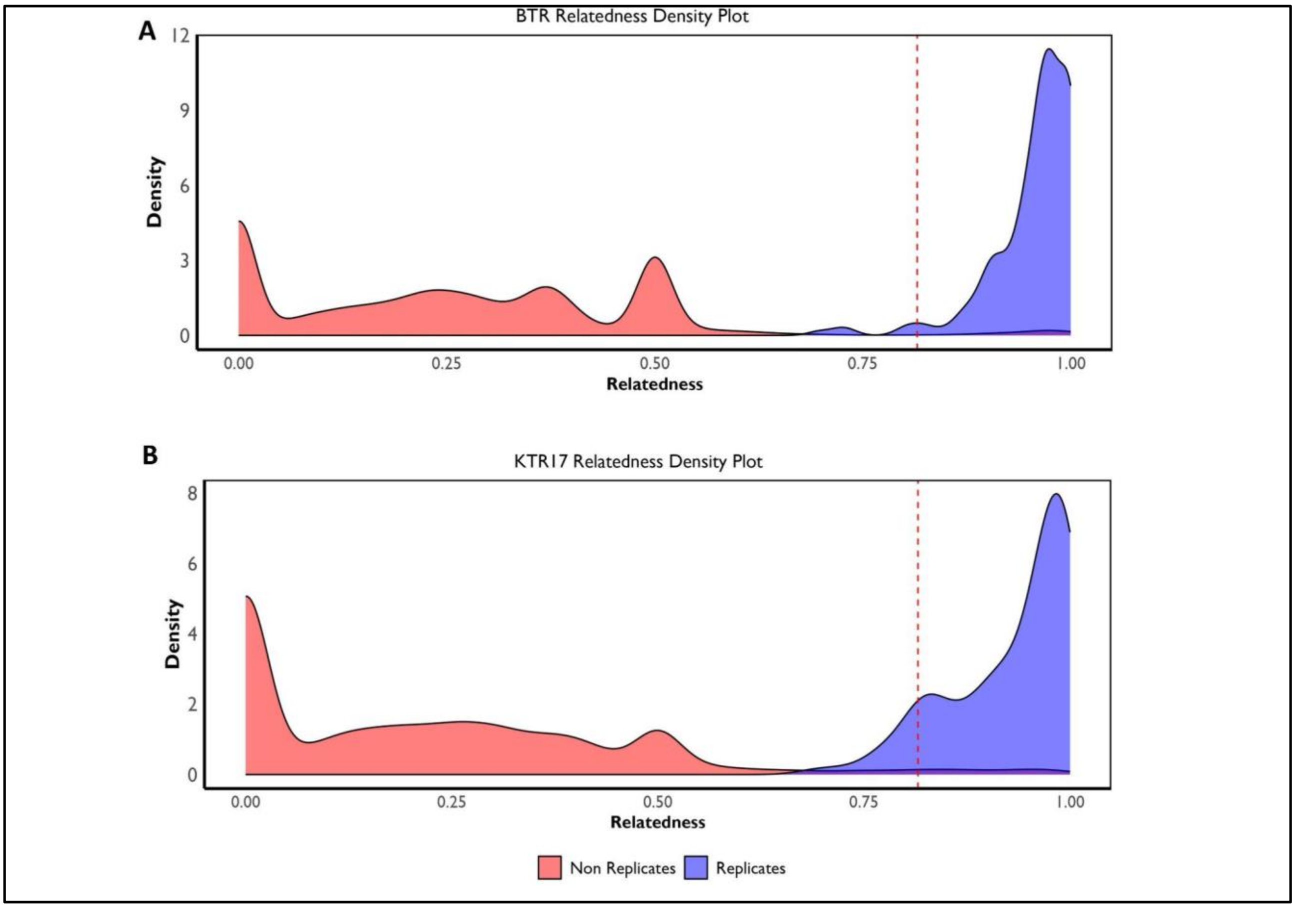


**Figure S2:** Relatedness density maps of Bandhavgarh (A) and Kanha (B). Recapture cut-off of 0.81 was set for both Kanha and Bandhavgarh based on 97.5% percentile of the relatedness distribution.

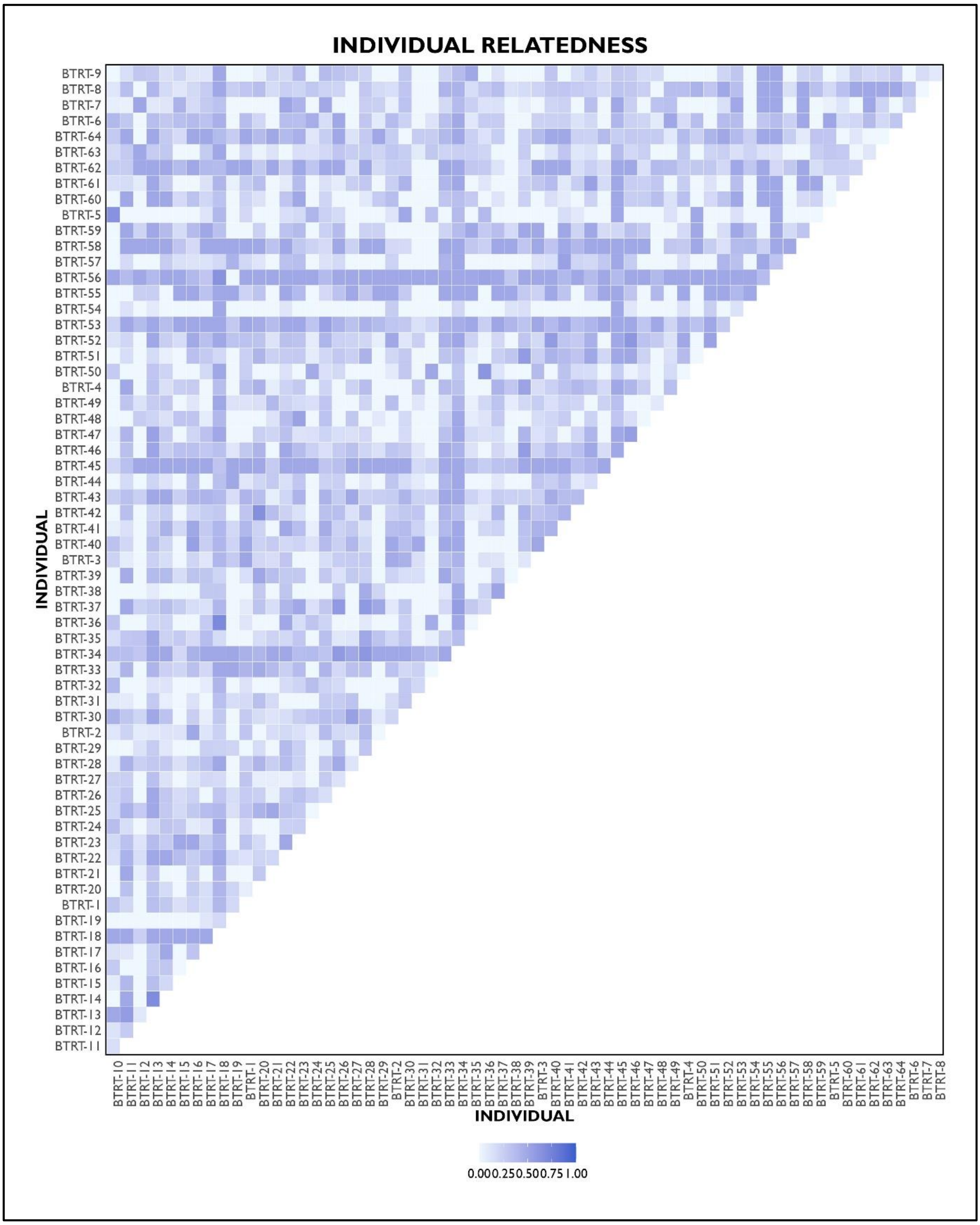


**Figure S3:** Pairwise relatedness estimated using PLINK between all pairs of individuals in Bandhavgarh.

.

| SEASON | SAMPLE TYPE | Probability of successful identification (mean) | SAMPLES (0.9) |
|---|---|---|---|
| Summer | SALIVA | 0.3 | 5.6 |
| Winter | SALIVA | 0.4 | 4.2 |
| Monsoon | SALIVA | 0.1 | 15.5 |
| Summer | SHED HAIR | 0.6 | 2.6 |
| Winter | SHED HAIR | 0.7 | 1.8 |
| Monsoon | SHED HAIR | 0.5 | 3.5 |

**Table S2:** Number of samples required per kill to identify an individual (with probability of 0.9).
